# Engineering CAR T Cells for Hepatocellular Carcinoma Recurrence after Liver Transplantation

**DOI:** 10.64898/2026.06.30.735569

**Authors:** Lorenz Kocheise, Gabriel Bacil, Pavan Bhimalli, Mohamed-Reda Benmebarek, Dan Li, Patrick Huang, Chi Ma, Vinona Muralidaran, Jorge Hernandez-Felix, Joseph R. Bugliarelli, Raj Chari, Kylynda Bauer, Yuta Myojin, Sarah Firdaus, Xiao Bin Zhu, Corbin Morris, Firouzeh Korangy, Alexander Kroemer, Mitchell Ho, Tim F. Greten

## Abstract

**Background & Aims:** Liver transplantation improves outcomes in hepatocellular carcinoma (HCC), yet treatment options for patients with tumor recurrence remain limited to tyrosine kinase inhibitors. Glypican-3 (GPC3)-targeted CAR T cells offer a tumor-directed immune-based therapeutic strategy, but their efficacy may be limited by post-transplant immunosuppression. We developed a CAR T cell platform combining CRISPR/Cas9-mediated *FKBP1A* disruption to confer resistance to FKBP12-dependent immunosuppressive agents, including tacrolimus, everolimus, and sirolimus, with *TRAC* knockout to eliminate endogenous T cell receptor expression and reduce alloreactivity.

**Methods:** Human T cells were edited using Cas9 ribonucleoprotein complexes targeting *FKBP1A* and *TRAC*, expanded, and transduced with an anti-GPC3 CAR construct. Cytokine production and cytotoxicity were assessed *in vitro*. Antitumor activity under tacrolimus treatment was evaluated in a Hep G2 xenograft model, and xenoreactivity was assessed in a graft-versus-host disease model. *FKBP1A/TRAC* double-knockout T cells were enriched using mTOR inhibitor selection combined with CD3-based MACS depletion. PBMCs from liver transplant recipients were used to evaluate feasibility for clinical translation during the early post-transplant period.

**Results:** Tacrolimus suppressed wild-type CAR T cell function but not *FKBP1A*/*TRAC* double-knockout CAR T cells, which retained cytokine production, cytotoxicity, and *in vivo* antitumor activity. Cyclosporine A remained suppressive, enabling its potential use as a pharmacologic control strategy. *TRAC* disruption reduced xenoreactivity. CD3-based MACS depletion and mTOR inhibition achieved functional double-knockout efficiencies greater than 98%, without compromising cell viability. Functional *FKBP1A/TRAC* knockout CAR T cells were generated from patient PBMC samples 30 days post-transplant.

**Conclusions:** Dual-edited GPC3 CAR T cells resist tacrolimus-based immunosuppression while limiting alloreactivity, supporting their use for recurrent HCC after liver transplantation. Sequential, high-viability selection in a modular cellular engineering framework enables adaptation to alternative tumor targets and next-generation CAR T cell designs.

**Impact and implications:**

- Engineering armored GPC3-targeting CAR T cells for hepatocellular carcinoma in the post–transplant setting
- Demonstrating safety and therapeutic efficacy in the presence of immunosuppressive drugs in preclinical models
- Leveraging mTOR inhibitor resistance and CD3 destabilization enables high-efficiency, high-purity selection of genetically modified T cells
- A modular cellular engineering platform enables utilization for alternative tumor targets and next-generation CAR T cells

## Introduction

For patients with resectable hepatocellular carcinoma (HCC), liver transplantation offers the best prospects for durable disease control [1, 2]. Nonetheless, post-transplant recurrence remains a major driver of cancer-related mortality, even among carefully selected candidates [3, 4]. Immune checkpoint inhibitors and other immunotherapy-based regimens carry a high risk of causing acute allograft rejection and are not used routinely after transplantation [1, 5, 6]. As a result, less effective tyrosine kinase inhibitor-based strategies remain the default systemic therapy for transplant recipients with recurrent disease [1]. Although transplant is typically reserved for curative intent in early-stage disease, selected patients with more advanced tumors and underlying end-stage liver disease may also derive substantial clinical benefit [7]. Given the persistent shortage of donor organs, candidates with borderline risk profiles may be deprioritized despite the possibility of meaningful benefit. Strategies that reduce recurrence and enable effective treatment in the immunosuppressed post-transplant setting could therefore expand eligibility criteria and allow more patients to benefit from transplantation.

Cellular immunotherapies, including chimeric antigen receptor (CAR) T cells, offer a tumor-directed approach that could address this unmet need [8]. Glypican-3 (GPC3) is a leading HCC target because it is highly expressed in many tumors and largely absent from healthy liver tissue, and early clinical studies of next-generation GPC3-directed CAR T cells have reported antitumor activity with limited off-tumor toxicity [9, 10]. However, liver transplant recipients require lifelong immunosuppression, which may blunt CAR T cell expansion and effector function *in vivo*. Conversely, engineering CAR T cells to resist immunosuppressive drugs raises a distinct concern: drug-resistant T cells could retain or amplify alloimmune responses and thereby increase the risk of graft injury if endogenous T cell receptor (TCR) activity is preserved [5].

Here, we describe the development of a GPC3-specific T cell product engineered to withstand the effects of commonly used post-transplant immunosuppressive agents, such as tacrolimus and everolimus, via CRISPR-mediated *FKBP1A* knockout. In addition, simultaneous knockout of the TCR eliminates the potential for graft rejection mediated by endogenous TCRs in these drug-resistant T cells.

## Materials and Methods

### Generation of CRISPR-edited CAR T cells

PBMCs from healthy volunteers were isolated by density-gradient centrifugation, followed by T cell enrichment. T cells were rested overnight in IL-7–supplemented T cell medium before CRISPR editing. sgRNAs targeting the *TRAC* and *FKBP1A* loci were complexed with Cas9 nuclease and delivered into T cells by electroporation. Following electroporation, cells were recovered in cytokine-supplemented medium containing IL-2, IL-7, and IL-15 and stimulated with CD3/CD28 beads or TransAct reagent. After 48 hours, T cells were transduced with a second-generation lentiviral vector encoding an anti-GPC3 CAR, as previously described [11]. Cells were expanded for an additional 5 to 7 days before use in functional *in vitro* and *in vivo* assays.

### Ethics statement

PBMCs were collected 30 days after liver transplantation from patients undergoing transplantation for HCC at MedStar Georgetown University Hospital. All patients provided informed consent for the collection of clinical information and tissue acquisition under an Institutional Review Board-approved protocol (IRB #2017-0365) and a material transfer agreement between Georgetown University Hospital and the U.S. National Cancer Institute (MTA #43655-18).

### Statistical analysis

Sample sizes for animal studies were determined based on prior experiments in our laboratory using similar or identical liver tumor models. For all models, animals were randomized and investigators were blinded to group allocation. Statistical analyses were performed using GraphPad Prism 10 (GraphPad Software). Differences between groups were assessed using unpaired Student’s t tests, one-way ANOVA, or two-way ANOVA, as appropriate. Data are shown as mean ± SD unless otherwise indicated. A p value < 0.05 was considered statistically significant. ns, not significant; *p < 0.05; **p < 0.01; ***p < 0.001; ****p < 0.0001.

## Results

### Post-transplant HCC recurrence remains a clinical challenge

First, we studied the potential clinical need for an adoptive T cell approach in patients with HCC recurrence after liver transplantation. To define recurrence rates under stringent transplant selection criteria, we reviewed large multicenter studies reporting post-transplant HCC recurrence in patients meeting established eligibility standards, including the Milan criteria (**Supp. Tab. 1**). Across these studies, recurrence rates ranged from 6.9% to 16.7%, depending on the duration of post-transplant follow-up.

To address this unmet need, we developed an armored T cell therapy platform based on a GPC3-specific chimeric antigen receptor (CAR) that is currently being evaluated clinically (NCT05003895). For liver transplant recipients requiring chronic immunosuppression, we disrupted *FKBP1A*, encoding FKBP12, to confer resistance to FKBP12-dependent agents including tacrolimus, sirolimus, and everolimus. We also disrupted the *TRAC* locus to reduce the risk of graft-directed alloreactivity mediated by the endogenous T cell receptor (TCR) in immunosuppression-resistant CAR T cells (**Fig. 1A**). Resting CD3-positive T cells were isolated from donor peripheral blood mononuclear cells (PBMCs) by magnetic-activated cell sorting (MACS) and electroporated with ribonucleoprotein (RNP) complexes consisting of high-fidelity Cas9 protein and single-guide RNAs (sgRNAs) targeting *FKBP1A* and *TRAC*.

**Fig. 1.**
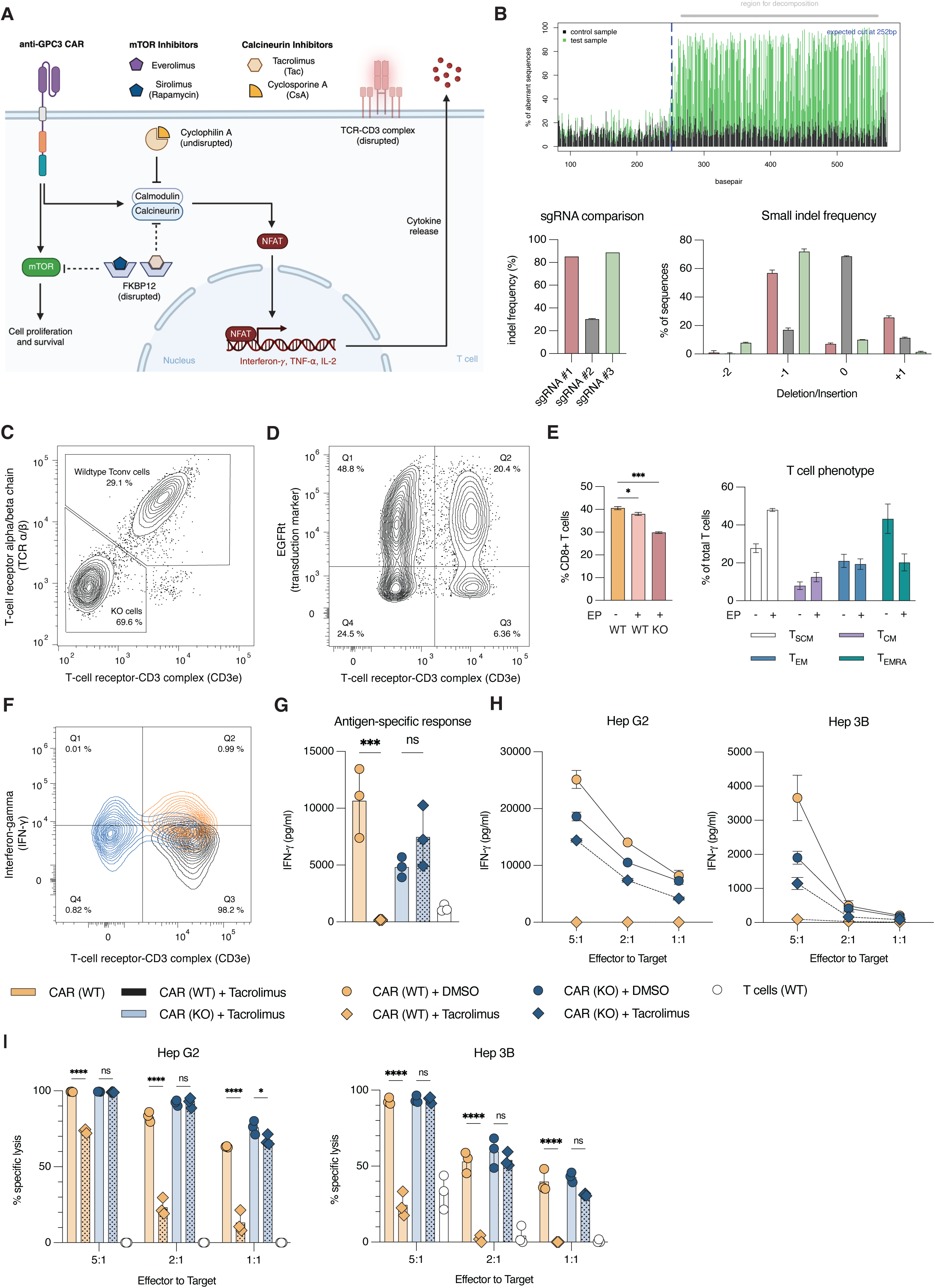
Genetic editing of anti-GPC3 CAR T cells and in vitro functional characterization. **(A)** Schematic of dual *FKBP1A*/*TRAC* disruption in second-generation anti-GPC3 CAR T cells. *FKBP1A* knockout prevents FKBP12-dependent inhibition by tacrolimus, sirolimus, and everolimus, while *TRAC* knockout reduces endogenous TCR-CD3 expression and TCR-mediated alloreactivity. Cyclosporine A remains active through cyclophilin A. **(B)** FKBP1A editing after CRISPR/Cas9 RNP electroporation, including representative TIDE decomposition, comparison of three sgRNAs, and indel distribution 7 days post-electroporation. **(C)** Representative flow cytometry plot showing surface TCRα/β and CD3ε expression 7 days after electroporation. TCRα/β-CD3ε double-negative cells were defined as TCR-deficient cells. **(D)** Representative flow cytometry of CAR transduction based on EGFRt, linked to the anti-GPC3 CAR by a 2A sequence, 7 days post-electroporation and 5 days post-transduction. **(E)** Frequency of CD8-positive cells among total T cells and T cell differentiation phenotype after expansion of non-electroporated and electroporated WT or KO T cells. T cell subsets were defined using CD45RA, CCR7, and CD95 staining, including stem cell memory-like T cells, central memory T cells, effector memory T cells, and terminally differentiated effector memory RA-positive T cells. **(F)** Intracellular IFN-γ staining after 4-hour PMA/ionomycin stimulation with brefeldin A in cells cultured with tacrolimus (15 ng/mL) or DMSO. **(G)** IFN-γ secretion after 48-hour culture of 1 × 10 CAR T cells on recombinant human GPC3-coated plates with tacrolimus or DMSO, measured by ELISA. **(H)** IFN-γ secretion and **(I)** luciferase-based cytotoxicity after 48-hour co-culture with 2 × 10 Hep G2-Luc or Hep 3B-Luc cells at indicated effector-to-target ratios with tacrolimus or DMSO. Graphs show representative experiments. Statistical significance was assessed by one-way or two-way ANOVA.

### Generation of FKBP12/TCR-deficient CAR T cells

We first evaluated whether disruption of *FKBP1A* could render engineered T cells resistant to FKBP12-dependent immunosuppressive agents, including tacrolimus, sirolimus, and everolimus. Multiple sgRNAs targeting *FKBP1A* were evaluated by Sanger sequencing, and editing efficiency was quantified using the TIDE algorithm based on forward and reverse sequencing reads (**Fig. 1B**) [12]. sgRNAs #1 and #3 produced the highest editing efficiencies, and sgRNA #1 was selected for subsequent experiments. Co-electroporation of sgRNAs targeting *FKBP1A* and *TRAC* generated double-knockout (KO) T cells, which were analyzed by flow cytometry for surface TCR expression (**Fig. 1C**). Disruption of *TRAC* eliminated TCRα/β and CD3ε surface staining, consistent with prior reports showing that stable surface CD3 expression depends on intact TCR expression. CD3 expression was therefore used as a surrogate marker for functional TCR deficiency [13].

Following electroporation, resting T cells were activated with CD3/CD28 beads and expanded for seven days in the presence of IL-2, IL-7, and IL-15. Two days after electroporation, cells were transduced with a second-generation lentiviral vector encoding a GPC3-specific CAR containing intracellular 4-1BB and CD3ζ signaling domains [11]. Seven days after activation, transduction efficiency was comparable between genetically modified and unmodified T cells (**Fig. 1D**). Electroporation caused a modest reduction in the percentage of CD8 cells, with a more pronounced effect in TCR-knockout cells (**Fig. 1E**). Electroporation was also associated with relative enrichment of T cells with a stem cell memory-like phenotype, defined by CD45RA, CCR7, and CD95 expression (**Fig. 1E**). In contrast, non-electroporated T cells showed relative enrichment of effector memory RA-positive cells. No phenotypic differences were detected between wild-type (WT) and knockout (KO) cells within the electroporated group, indicating that the observed changes were attributable to the electroporation procedure rather than to the specific gene knockouts (data not shown).

### FKBP12/TCR-deficient CAR T cells resist tacrolimus-mediated suppression

Next, we assessed FKBP12 deficiency in vitro. Tacrolimus suppressed PMA/ionomycin-induced IFN-γ production in WT T cells but not in FKBP12/TCR double-knockout T cells (**Fig. 1F**). Similarly, after GPC3-specific stimulation with recombinant GPC3-IgG1 Fc, tacrolimus inhibited IFN-γ production by WT CAR T cells, whereas KO CAR T cells maintained cytokine production (**Fig. 1G**). In GPC3-positive Hep 3B and Hep G2 tumor co-cultures, tacrolimus impaired WT CAR T cytokine production and cytotoxicity but not KO CAR T cell function (**Fig. 1H** and **Fig. 1I**).

### FKBP12/TCR-deficient CAR T cells retain antitumor activity under tacrolimus treatment *in vivo*

After successful *in vitro* characterization, we assessed the impact of immunosuppressive therapy on CAR T cell-mediated HCC treatment *in vivo*. To model post-transplant HCC recurrence, NSG mice were injected intraperitoneally with 2 × 10^6^ luciferase-expressing Hep G2 cells and monitored for 14 days to allow establishment of tumor burden. Mice then received 2 × 10 CAR T cells, followed by tacrolimus administered at a total daily dose of 1 mg/kg for five consecutive days. Tumor burden was quantified by IVIS imaging (**Fig. 2A**). Tacrolimus alone did not affect tumor burden in mice that did not receive CAR T cell therapy (**Fig. 2B**). KO CAR T cells induced a reduction in tumor burden comparable to WT CAR T cells in the absence of tacrolimus (**Fig. 2C**). In tacrolimus-treated mice, however, WT CAR T cells failed to control tumor growth while immunosuppression was maintained, as shown at day 3. After tacrolimus withdrawal, tumor burden in WT CAR T cell-treated mice decreased relative to untreated controls, suggesting restoration of antitumor activity by persisting WT CAR T cells once immunosuppression ceased (**Fig. 2D** and **Fig. 2E**). By contrast, KO CAR T cells maintained antitumor activity during tacrolimus treatment, with tumor control comparable to that observed in vehicle-treated KO CAR T cell recipients.

**Fig. 2.**
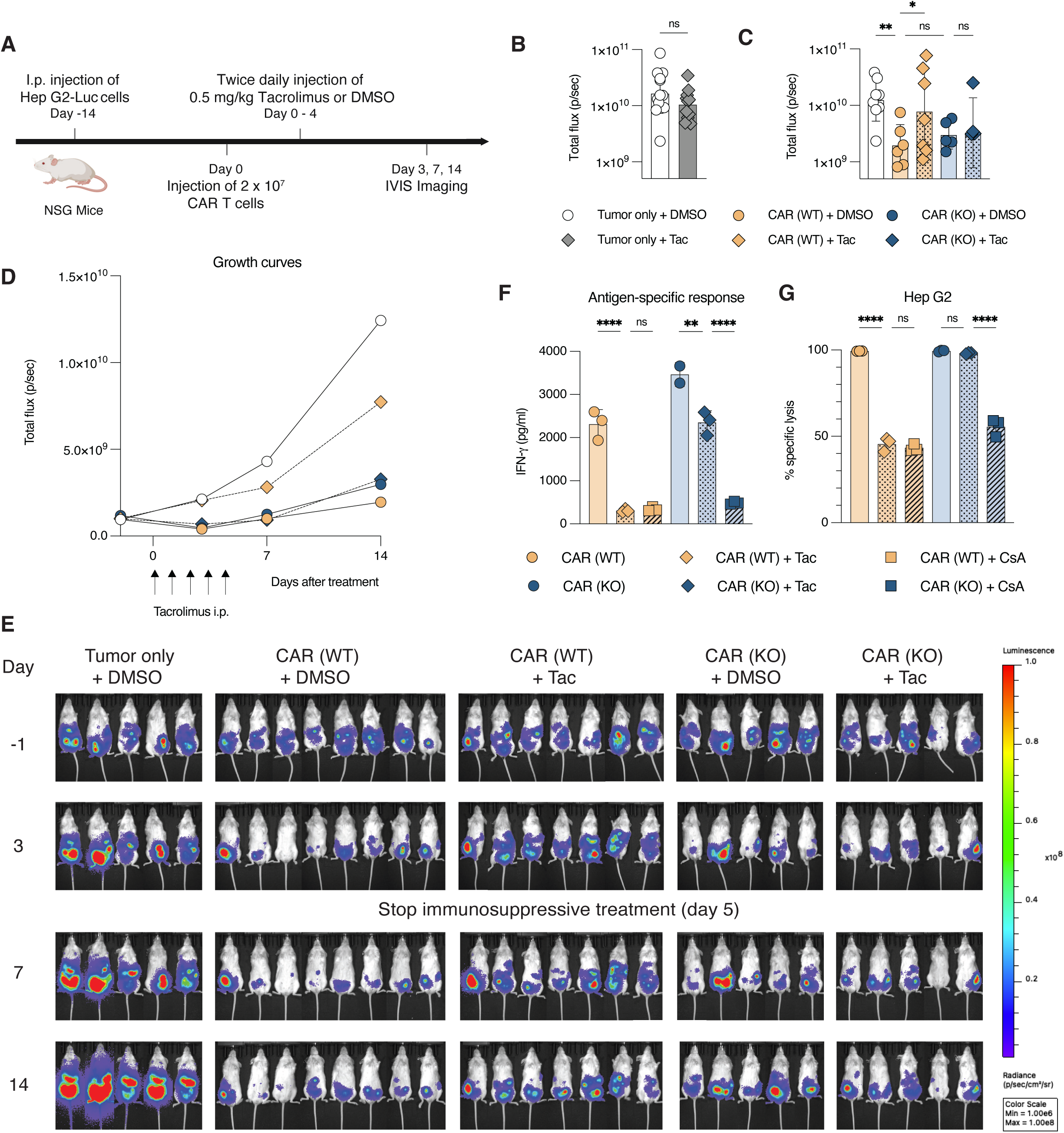
Armored CAR T cells retain antitumor activity during tacrolimus treatment in vivo and remain suppressible by cyclosporine A. **(A)** Schematic of the Hep G2 peritoneal tumor model. NSG mice received 2 × 10 Hep G2-Luc cells intraperitoneally on day -14, were randomized on day 0, and treated with 2 × 10 WT or KO CAR T cells. Mice received DMSO or tacrolimus, starting at 1 mg/kg followed by 0.5 mg/kg intraperitoneally twice daily for 4 days. Tumor progression was monitored by IVIS on days 3, 7, and 14. **(B)** Tumor burden in tumor-only mice treated with DMSO or tacrolimus, shown as pooled geometric mean ± SD from two experiments. **(C)** Total bioluminescent flux on day 14 in mice receiving WT or KO CAR T cells with DMSO or tacrolimus, shown as geometric mean ± SD from one representative experiment of two. **(D)** Tumor growth curves based on geometric mean IVIS signal; arrows indicate tacrolimus dosing. **(E)** Representative IVIS images before treatment and on days 3, 7, and 14 after CAR T cell infusion. **(F)** IFN-γ secretion after 48-hour culture of 1 × 10 CAR T cells on recombinant human GPC3-coated plates with tacrolimus (15 ng/mL), cyclosporine A (300 ng/mL), or vehicle control, measured by ELISA. **(G)** Luciferase-based cytotoxicity of WT and KO CAR T cells against Hep G2-Luc cells with tacrolimus, cyclosporine A, or vehicle control. Statistical significance was assessed by one-way or two-way ANOVA.

### Cyclosporine A provides pharmacologic control of FKBP12-deficient CAR T cells

Although FKBP12 disruption enables CAR T function during tacrolimus exposure, CAR T activity may still need to be controlled to manage toxicity. We therefore tested cyclosporine A, a calcineurin inhibitor that binds cyclophilin A rather than FKBP12. Tacrolimus and cyclosporine A comparably suppressed IFN-γ production and cytotoxicity in WT CAR T cells (**Fig. 2F** and **Fig. 2G**). KO CAR T cells remained sensitive to cyclosporine A, suggesting a clinically available strategy to suppress FKBP12-deficient CAR T cells if needed.

### *TRAC* disruption prevents xenoreactive liver injury *in vivo*

We next assessed the safety profile of this CAR T cell platform in the transplantation setting. After liver transplantation, the graft remains an allogeneic organ at ongoing risk of immune-mediated rejection, necessitating chronic immunosuppression. Although FKBP12 deficiency renders KO CAR T cells resistant to agents such as tacrolimus and everolimus, these armored T cells could theoretically promote graft rejection through their endogenous TCR. Disruption of *TRAC* was therefore incorporated to abrogate native TCR expression and restrict antigen recognition to CAR specificity. RNP electroporation targeting *FKBP1A* and *TRAC* achieved over 70% knockout efficiency, as indicated by loss of TCRα/β and CD3ε staining. However, a substantial fraction of T cells remained unmodified and retained endogenous TCR expression. To reduce this population, we implemented a purification strategy based on negative selection of CD3-positive cells, taking advantage of CD3 complex destabilization after TCR disruption (**Fig. 3A**). Conventional MACS-based depletion of CD3-positive cells yielded products with up to 99% CD3-negative purity without compromising cell viability.

**Fig. 3.**
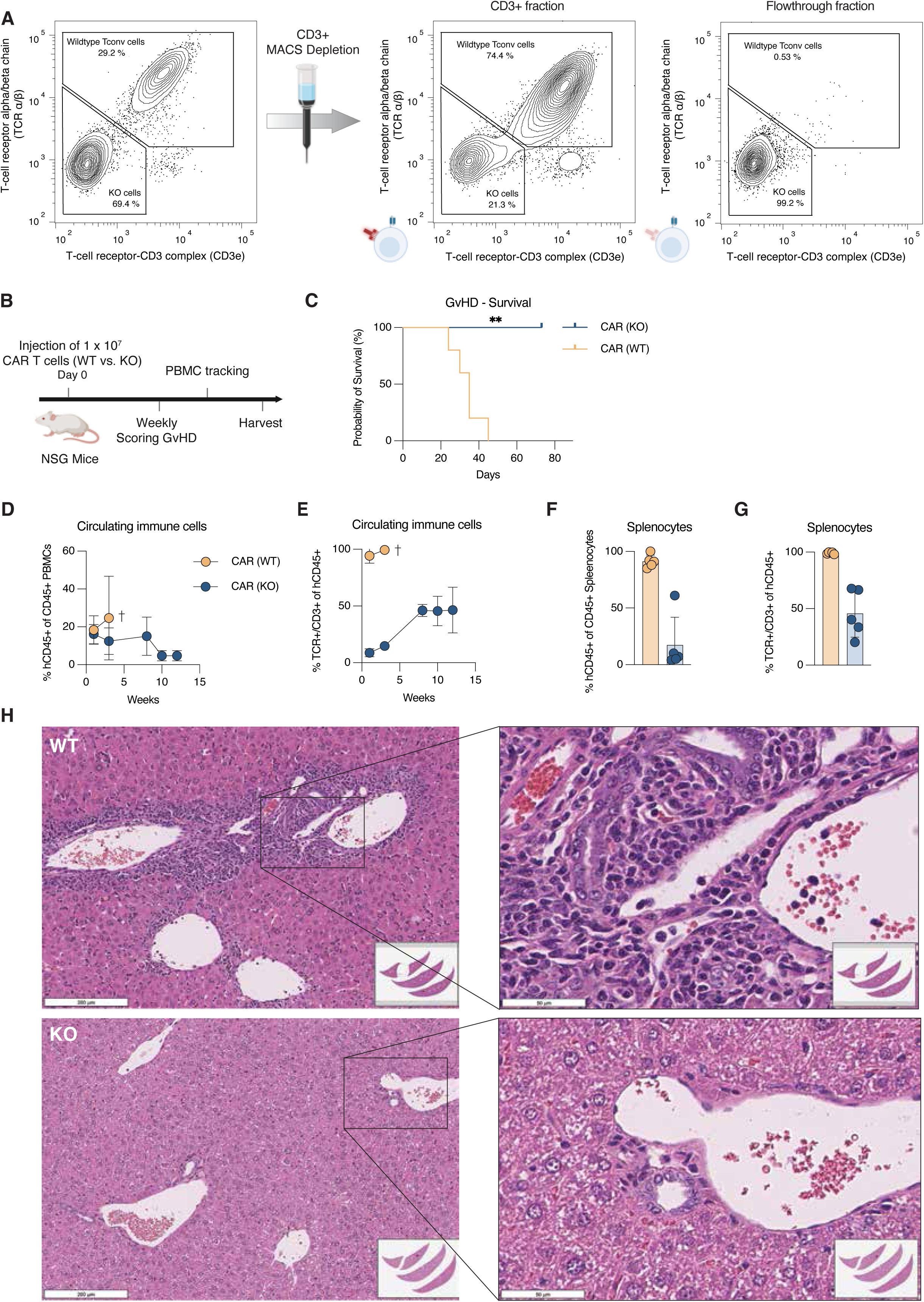
TRAC disruption and CD3 depletion reduce xenoreactive T cell expansion and liver injury in vivo. **(A)** Enrichment of TCR/CD3-negative KO T cells by CD3 depletion. Representative flow cytometry plots show TCRα/β and CD3ε expression before CD3 depletion, in the CD3-positive retained fraction, and in the CD3-negative flowthrough fraction. **(B)** Schematic of the in vivo safety experiment. NSG mice were irradiated with 1.3 Gy and injected intravenously with 1 × 10^7^ WT CAR (n = 5) T cells or CD3-depleted KO (n = 5) CAR T cells. Mice were monitored weekly for clinical signs of graft-versus-host disease, and peripheral blood was collected longitudinally to track circulating human T cells. Spleens and livers were harvested at euthanasia or at the experimental endpoint. **(C)** GvHD-dependent survival shown by Kaplan-Meier analysis. Survival curves were analyzed using the log-rank test. **(D)** Frequency of circulating human T cells over time, defined as the proportion of human CD45-positive cells among total (human + murine) CD45-positive peripheral blood cells. **(E)** Frequency of residual TCR/CD3-positive cells among circulating human CD45-positive cells over time. **(F)** Frequency of human CD45-positive cells among total CD45-positive splenocytes at euthanasia or experimental endpoint. **(G)** Frequency of residual TCR/CD3-positive cells among human CD45-positive splenocytes. **(H)** Representative H&E-stained liver sections from mice treated with WT or KO CAR T cells. Scale bars, 200 µm in low-magnification images and 50 µm in high-magnification images.

To evaluate alloreactive potential of the enriched KO CAR T cell products *in vivo*, we used a xenogeneic graft-versus-host disease (GvHD) model as a surrogate for graft-directed reactivity. NSG mice were irradiated with 1.3 Gy and injected with 1 × 10^7^ WT or KO CAR T cells (**Fig. 3B**). Mice receiving KO T cells showed no clinical signs of GvHD throughout the 89-day observation period. By contrast, mice treated with WT T cells rapidly developed severe clinical GvHD and were euthanized according to predefined criteria (**Fig. 3C**). In mice receiving WT T cells, circulating human T cell numbers increased steadily over time. In contrast, mice injected with KO T cells exhibited a progressive decline in circulating human T cell counts (**Fig. 3D**). Further analysis of the KO group revealed a gradual increase in the proportion of residual unmodified T cells over time (**Fig. 3E**), suggesting selective expansion of rare TCR-positive clones. Consistent with these findings, more than 90% of CD45-positive splenocytes in WT T cell-treated mice were of human T cell origin at the time of euthanasia (**Fig. 3F**). In the KO group, this proportion varied substantially, consistent with stochastic expansion of residual xenoreactive clones. Mirroring the expansion of unmodified T cells in peripheral blood, splenic human T cells from KO-treated mice showed an increased frequency of residual endogenous TCR expression compared with the initial injection product (**Fig. 3G**). These findings underscore the importance of minimizing residual endogenous TCR-positive cells in the final product to prevent clonal expansion of graft-reactive, immunosuppression-resistant T cell populations.

Histopathological evaluation showed no evidence of liver injury in mice treated with KO T cells. In contrast, livers from WT T cell-treated mice exhibited pronounced pathological alterations (**Fig. 3H**). Portal tracts were moderately to markedly expanded by mixed inflammatory infiltrates composed predominantly of lymphocytes, with fewer macrophages and neutrophils. These infiltrates were associated with hepatocellular necrosis or apoptosis. Inflammation also extended beneath the endothelium of portal and central veins, which displayed variably plump, reactive endothelial cells. The biliary epithelium was disorganized, with irregular spacing, epithelial overlap, mild nuclear atypia, and occasional lymphocytic infiltration. Multifocal dilation of lymphatic vessels was also observed.

### Pharmacologic and magnetic selection strategies enrich double-edited T cells

Removing TCR-expressing cells from the final knockout CAR T cell product and enriching for FKBP12-deficient CAR T cells would be desirable before clinical translation. We therefore investigated whether FKBP12 deficiency could be leveraged to enrich for edited T cells (**Fig. 4A**). Addition of the FKBP12-dependent mTOR inhibitors everolimus or sirolimus to T cell cultures beginning two days after electroporation and activation selectively enriched cells harboring edits at the *FKBP1A* locus (**Fig. 4B**). By contrast, the NFAT inhibitor tacrolimus did not enrich FKBP12-deficient cells, likely because exogenous cytokines in the culture reduced dependence on autocrine and paracrine activation signals. Enrichment of FKBP12-deficient cells was accompanied by increased indel frequencies at the *TRAC* locus (**Fig. 4C**). We hypothesized that a fraction of non-electroporated cells retained unmodified genomic loci at both *FKBP1A* and *TRAC*, thereby diluting the overall editing frequency. Notably, the indel frequency at the *TRAC* locus correlated with flow cytometry staining but did not match exactly, most likely due to allelic exclusion of the unmodified TRAC allele. Consistent with this interpretation, genomic sequencing of CD3 MACS-sorted T cells showed that depletion of CD3-positive cells (**Fig. 4D**), which enriches for TCR-knockout cells (**Fig. 4E**), also increased the fraction of *FKBP1A*-modified T cells (**Fig. 4F**).

**Fig. 4.**
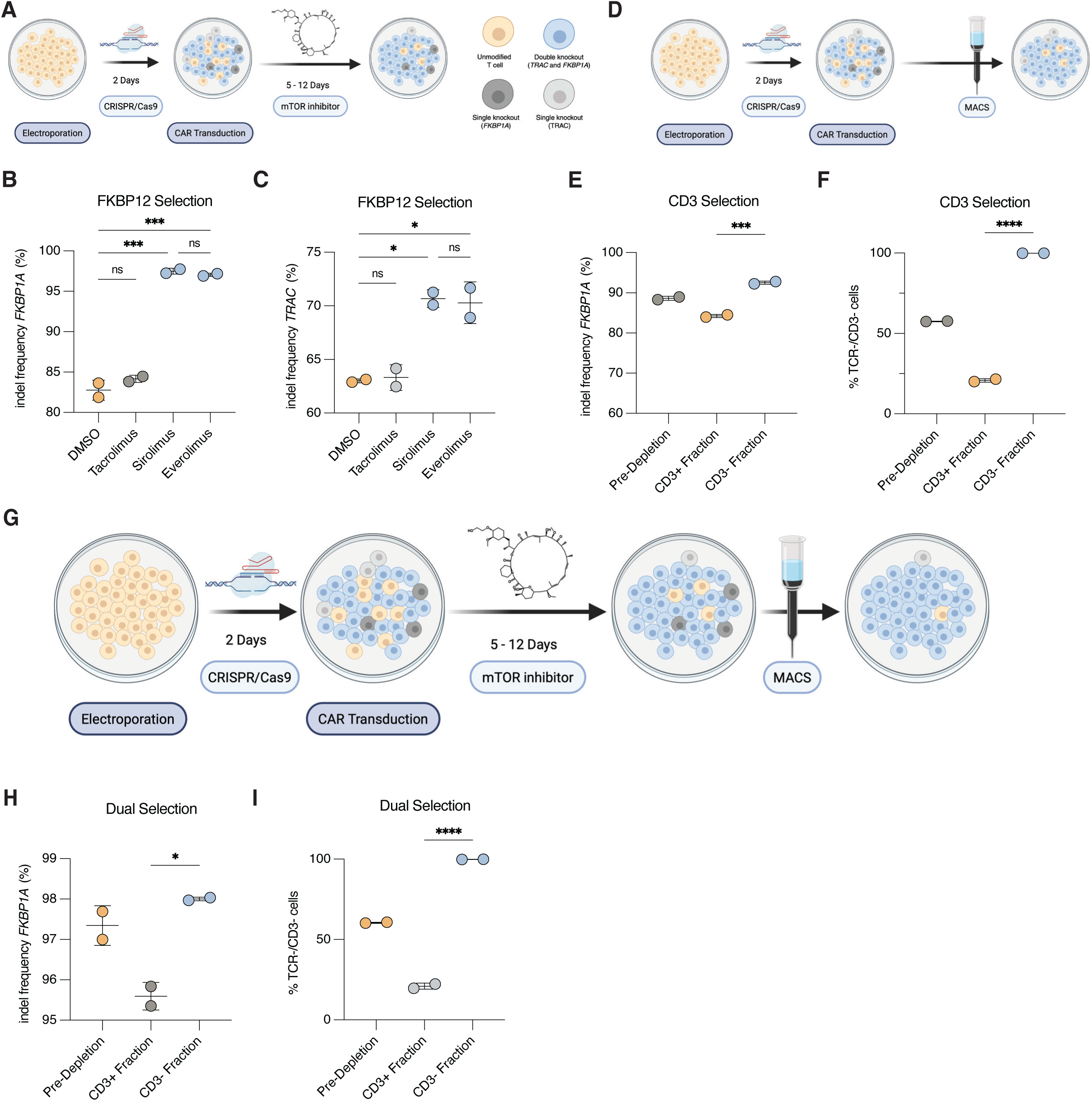
Sequential pharmacologic and magnetic enrichment yields highly pure, viable FKBP1A/TRAC double-knockout T cells. **(A)** Pharmacologic enrichment of *FKBP1A*-edited T cells during *ex vivo* expansion. Cells were cultured with DMSO (1:10,000), tacrolimus (100 ng/mL), sirolimus (100 ng/mL), or everolimus (100 ng/mL). **(B)** *FKBP1A* indel frequencies were quantified at the end of expansion. **(C)** *TRAC* indel frequencies after pharmacologic selection with DMSO (1:10,000), tacrolimus (100 ng/mL), sirolimus (100 ng/mL), or everolimus (100 ng/mL). **(D)** MACS-based enrichment of TCR/CD3-negative cells after CRISPR/Cas9 editing. **(E)** Frequency of TCRα/β-CD3ε double-negative cells is shown before CD3 depletion, in the CD3-positive retained fraction, and in the CD3-negative untouched flowthrough fraction. **(F)** *FKBP1A* indel frequency after CD3-based magnetic selection, assessed by genomic NGS sequencing. **(G)** Schematic of the combined enrichment strategy. T cells were electroporated with CRISPR/Cas9 RNPs targeting *FKBP1A* and *TRAC*, expanded in the presence of an mTOR inhibitor to enrich *FKBP1A*-disrupted cells, and then subjected to CD3-positive MACS depletion to remove residual TCR/CD3-positive cells. **(H)** *FKBP1A* indel frequencies and **(I)** frequencies of TCRα/β-CD3ε double-negative cells after combined pharmacologic FKBP12 selection and CD3-based magnetic depletion. Statistical significance was assessed using one-way ANOVA.

Because pharmacologic and magnetic selection enrich complementary features of the edited product, we next investigated whether both strategies could be combined within a single manufacturing workflow (**Fig. 4G**). Everolimus or sirolimus treatment was used to enrich *FKBP1A*-disrupted cells during expansion, whereas subsequent CD3-positive depletion removed residual TCR/CD3-positive cells and increased the purity of the TCR-negative fraction (**Fig. 4F**). The combined selection strategy resulted in high frequencies of double-edited cells, yielding *FKBP1A* modification rates of 98.0% and a TCR/CD3-negative-cell frequency of 99.9% (**Fig. 4H** and **Fig. 4I**).

### Gene-editing platform enables alternative HCC antigen targeting and CAR T cell generation from post-transplant patient PBMCs

Although GPC3 is expressed in approximately 70% of HCC samples, we also explored whether this gene-editing platform could be applied to alternative HCC targets [14]. To demonstrate the versatility of our approach, we used a previously described HLA-A2-restricted alpha-fetoprotein (AFP)-specific TCR [15]. T cells were transduced on day 2 after activation using a second-generation lentiviral system to deliver the AFP-TCR in place of the GPC3-CAR. We demonstrated that KO T cells retained antigen-specific cytotoxicity in the presence of tacrolimus. In contrast, KO T cells remained susceptible to cyclosporine A, consistent with our observations in GPC3-targeting CAR T cells (**Fig. 5A** and **Fig. 5B**).

**Fig. 5.**
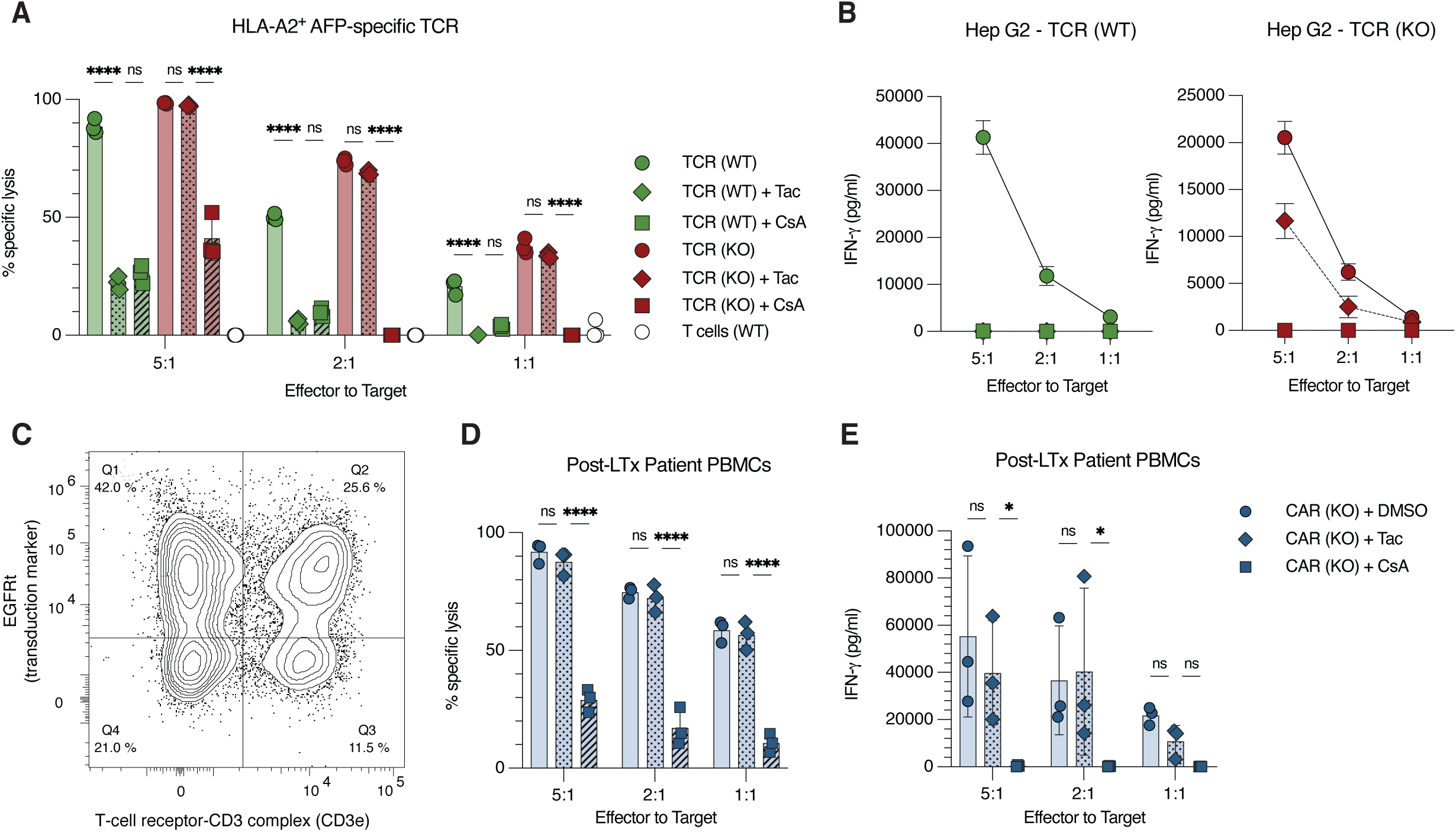
Modular engineering enables versatile cell therapy design and clinical translation. **(A)** Luciferase-based cytotoxicity of WT and KO HLA-A2-restricted AFP-specific TCR T cells against HLA-A2-positive Hep G2-Luc cells at indicated effector-to-target ratios with tacrolimus (15 ng/mL), cyclosporine A (300 ng/mL), or DMSO. **(B)** IFN-γ secretion by AFP-specific TCR T cells after Hep G2 co-culture under the same conditions. **(C)** Representative flow cytometry of EGFRt and CD3ε expression in edited and expanded CAR T cells generated from post-liver-transplant patient PBMCs; one representative of five patient-derived samples is shown. **(D)** Luciferase-based cytotoxicity of patient PBMC-derived KO CAR T cells against Hep G2-Luc cells with DMSO, tacrolimus, or cyclosporine A. Each data point represents the average cytotoxic activity from one of three patients. **(E)** IFN-γ secretion by patient PBMC-derived KO CAR T cells after Hep G2 co-culture with DMSO, tacrolimus, or cyclosporine A. Each data point represents the average IFN-γ production from one of three patients. Statistical significance was assessed by two-way ANOVA.

To assess the clinical relevance of our approach, we analyzed PBMCs from patients 30 days after liver transplantation to evaluate their suitability for generating a clinically relevant cell product. Following successful *ex vivo* expansion, electroporation, and CAR transduction, patient-derived PBMCs generated CAR T cells with cytotoxic activity and cytokine production comparable to healthy donor controls (**Fig. 5C**, **Fig. 5D**, and **Fig. 5E**). Consistent with our prior findings, these CAR T cells demonstrated resistance to tacrolimus while remaining sensitive to cyclosporine A, reinforcing the clinical applicability of our approach. Collectively, these data indicate that PBMCs collected as early as 30 days after liver transplantation can serve as a viable source for generating FKBP12/TCR-deficient GPC3-targeting CAR T cells, supporting future clinical translation of this approach.

## Discussion

We developed a GPC3-directed CAR T cell platform incorporating dual disruption of *FKBP1A*, which encodes FKBP12, and *TRAC*, which encodes the T cell receptor alpha chain, to enable immunotherapy for HCC recurrence after liver transplantation. This strategy was designed to preserve CAR T cell antitumor activity under immunosuppression while reducing TCR-mediated recognition of allogeneic liver tissue. We achieved high dual-knockout efficiencies using electroporation. Residual TCR-positive cells were further depleted by standard CD3 MACS selection, increasing TCR-knockout purity to as high as 99%. In the absence of immunosuppressive drugs, wild-type and knockout CAR T cells showed comparable cytotoxicity and cytokine production, indicating that *FKBP1A* and *TRAC* disruption did not substantially impair baseline CAR T cell function. Importantly, *FKBP1A* disruption conferred resistance to tacrolimus by preventing tacrolimus-mediated inhibition of calcineurin/NFAT signaling, which we confirmed both *in vitro* and *in vivo*. Because *FKBP1A* knockout also renders CAR T cells resistant to the mTOR inhibitors sirolimus and everolimus, while cyclosporine A signaling through cyclophilin remains intact, this platform provides a clinically useful pharmacologic control strategy. Tacrolimus or mTOR inhibitors could be used to selectively suppress the host immune compartment while preserving CAR T cell activity, whereas cyclosporine A could be used to suppress the entire immune compartment, including the engineered CAR T cells, in the event of unexpected CAR-mediated toxicity.

Simultaneous disruption of the endogenous TCR provides an additional safety layer by reducing both direct recognition of donor HLA molecules and indirect recognition of donor-derived antigens presented by host antigen-presenting cells. In a xenograft GvHD model, *TRAC* disruption protected against HLA-mediated alloreactivity. Notably, even partially edited products containing substantial numbers of unmodified T cells showed protection, consistent with the concept that reducing the number of functional endogenous TCRs decreases the likelihood of alloreactive clonal expansion. Together, these findings support dual *FKBP1A*/*TRAC* knockout as a strategy to preserve CAR-mediated antitumor function while limiting both pharmacologic suppression and TCR-mediated graft injury.

Sirolimus and everolimus mediate mTOR inhibition through FKBP12 and are widely used for immunosuppression after liver transplantation [16]. mTOR inhibitors may be especially suitable for post-transplant HCC recurrence, because of their potential antitumor effects [17]. We leveraged the resistance of *FKBP1A*-knockout cells to these drugs to selectively enrich *FKBP1A*-disrupted T cells. Interestingly, this approach also enriched for TCR-knockout cells, suggesting preferential co-delivery or uptake of both sgRNAs within the edited T cell population. By combining mTOR inhibitor-based selection with CD3-positive MACS depletion, we established a practical and low-cost enrichment strategy that preserves T cell viability while yielding an optimized cell product. Because CD3 depletion is a negative-selection step and *FKBP1A*-knockout cells are resistant to mTOR inhibitor signaling, this approach is expected to be compatible with clinical translation.

CRISPR/Cas9-modified T cell products have been proposed for the treatment of viral infections and hematologic malignancies after stem cell transplantation [18–20]. HCC is the only solid cancer for which organ transplantation represents a routinely used treatment approach [21]. To our knowledge, this is the first study to combine *FKBP1A*/*TRAC* editing with GPC3-directed CAR T cells for HCC recurrence after liver transplantation. Previously, the Bertoletti group pioneered cellular therapies to address HCC recurrence after liver transplantation. By electroporating anti-HBV TCR mRNA, they achieved transient expression of TCRs targeting hepatitis B-positive HCCs and co-electroporated mRNA encoding drug-resistant variants of calcineurin B (CnB) and inosine-5′-monophosphate dehydrogenase (IMPDH) to temporarily protect these cells from immunosuppressive drugs [22]. While this approach is intriguing for hepatitis-positive HCC patients, it also limits the durability of receptor expression and pharmacologic resistance, making it challenging to implement the robust, stable genetic engineering and armoring of CAR T cells that will likely be required for future approaches.

Recent early clinical trials using next-generation armored CAR T cell products have renewed optimism regarding the potential of CAR T cell therapy for solid tumors [8, 9, 23]. A key advantage of our platform is that the CRISPR/Cas9 editing module is independent of lentiviral CAR transduction and can therefore be adapted to a range of receptor designs. For example, the safety of an AFP-targeted TCR T cell therapy was recently demonstrated in a phase I study and could be incorporated into this manufacturing platform [24]. *HLA-A*02:01-restricted AFP-targeted TCRs may be particularly attractive in the setting of HLA-mismatched liver transplantation, as they could selectively recognize AFP-expressing recipient-derived HCC cells while sparing HLA-mismatched donor hepatocytes. By lentiviral transduction of double-knockout T cells with a previously described *HLA-A*02:01*-*restricted AFP-targeting TCR, we demonstrated the versatility of this approach for incorporating diverse genetic constructs [15]. Similarly, hepatitis virus-specific TCRs could be stably incorporated into this platform [25]. Although treatment after HCC recurrence represents the most immediate clinical application, these armored CAR T cells could also be manufactured before liver transplantation and administered early after transplant to target minimal residual disease. Treating patients at a lower tumor burden may increase the likelihood of complete tumor eradication, particularly given the emergence of antigen escape as an important resistance mechanism in GPC3-directed CAR T cell therapy.

Recent work from the Carl June laboratory demonstrated that dual-targeted CD19 and mesothelin CAR T cells can improve treatment of solid tumors by using peripheral B cell depletion to sustain CAR T cell persistence, reduce the need for lymphodepletion, and improve tumor clearance [26]. A similar strategy could be considered for immunosuppression-resistant armored CAR T cells in the transplant setting. In this context, B cell depletion might not only enhance CAR T cell activity but also contribute to suppression of humoral alloimmune responses in liver transplant recipients.

Beyond HCC, this strategy may also be relevant for post-transplant lymphoproliferative disorder (PTLD), for which treatment approaches with CD19-targeted CAR T cells have been explored [27]. A platform that allows continuation of high-dose immunosuppression while preserving cellular therapy efficacy could be particularly valuable in this setting. The importance of this concept was highlighted by a recent case report describing acute allogeneic rejection following CAR T cell treatment for PTLD after liver transplantation [28].

Several limitations must be addressed before clinical translation. Xenograft tumor models only partially capture the immunologic complexity of liver transplantation, including liver-specific tolerance, chronic pharmacologic immunosuppression, donor-specific alloimmunity, and tumor-associated immune suppression. Further studies are also needed to evaluate long-term persistence and chromosomal rearrangements of genetically edited CAR T cells. Importantly, site-specific integration strategies, such as SEED-mediated insertion of genetic cargo into intronic regions of *FKBP1A* and *TRAC*, may enable more uniform transgene expression while linking therapeutic cargo delivery to successful knockout [29].

Expanding eligibility criteria for liver transplantation is expected to increase rates of hepatocellular carcinoma recurrence in the coming years [30]. At the same time, rapid progress in xenotransplantation presents new opportunities and challenges for patients with difficult-to-treat HCC [31]. The reported transplantation of a genetically modified swine liver into a patient with HCC and impaired liver function represents an important milestone and highlights the potential relevance of xenotransplantation in this population [32]. A new generation of armored CAR T cells designed to function despite intense immunosuppression could help realize the therapeutic potential of both allogeneic and xenogeneic transplantation for patients with HCC.

## Abbreviations

AFP: alpha-fetoprotein
CAR: chimeric antigen receptor
GPC3: glypican-3
GvHD: graft-versus-host disease
HCC: hepatocellular carcinoma
IFN-γ: interferon gamma
KO: knockout
MACS: magnetic-activated cell sorting
NSG: NOD-*scid* IL-2Rg^null^ mice
PBMCs: peripheral blood mononuclear cells
sgRNA: single-guide RNA
TCR: T cell receptor
WT: wild type.

## Acknowledgement

This research was supported in part by the Intramural Research Program of the National Institutes of Health (NIH). The contributions of the NIH authors are considered Works of the United States Government. The findings and conclusions presented in this paper are those of the authors and do not necessarily reflect the views of the NIH or the U.S. Department of Health and Human Services.

## Declaration of generative AI use

During the preparation of this manuscript, the authors used GPT-5.5 Pro (OpenAI) to improve the readability and clarity of the text. The authors reviewed and edited the results and take full responsibility for the content of the published article.

## Author contributions

Conceptualization: LK and TFG; Formal analysis: LK, GB, PH, JHF, JRB, RC, and TFG; Investigation: LK, GB, PB, MRB, DL, PH, CM, VM, JRB, RC, JHF, KB, YM, XBZ, SF, CoM, FK, AK, MH, and TFG; Resources: LK, DL, VM, AK, MH, and TFG; Writing - Original Draft: LK and TFG; Writing - Review & Editing: LK, GB, PB, MRB, DL, PH, CM, VM, JRB, RC, JHF, KB, YM, XBZ, SF, CoM, FK, AK, MH, and TFG; Visualization: LK and TFG.

## Data availability statement

Data generated during this study will be made available from the corresponding author upon request.

