## Supplementary Table 1 for "Engineering CAR T Cells for Hepatocellular Carcinoma Recurrence after Liver Transplantation"

Tables

**Supplementary Table 1.** **Contemporary multicenter cohorts reporting HCC recurrence after liver transplantation**

| **Region and study** | **Study years** | **n (transplanted)** | **Selection criteria** | **Recurrence endpoint** |
| --- | --- | --- | --- | --- |
| U.S.; Mehta et al., 2020  (Hepatology)  PMID: 31344273 | 2012–2015 | 3,819 | Within Milan criteria | 3‑year recurrence rate  6.9%. |
| U.S.; Mehta et al., 2020  (Hepatology)  PMID: 31344273 | 2012–2015 | UNOS-DS: 422;  AC-DS: 121 | Expanded criteria | 3‑year recurrence rate  12.8% (UNOS‑DS)  and 16.7% (AC‑DS). |
| U.S.; Agopian et al., 2017  (Ann Surg)  PMID: 28654545 | 2002–2013 | 3,601 | Within Milan criteria | 5-year recurrence rate  11.2% (bridged) vs 10.1% (no LRT). |
| U.S.; Tabrizian et al., 2022  (JAMA Surg)  PMID: 35857294 | 2001–2015 | 2,645 | Within Milan criteria | 10-year recurrence rate 13.3% |
| Brazil; Chagas et al., 2019  (Eur J Gastroenterol Hepatol)  PMID: 31247632 | 2006–2015 | 1,119 | Multiple criteria. | Overall recurrence rate  8% |
| Japan; Shimamura et al., 2019 (Transplant Int)  PMID: 30556935 | 1998–2009 | 965  (LDLT) | “5-5-500 rule” ( tumor size ≤5 cm, tumor number ≤5, AFP ≤500 ng/mL) | 5-year recurrence rate  7.3% |
| U.S.; Tran et al., 2023 (Liver Transpl)  PMID: 37029083 | 2002–2013 | 4,981 | Multiple criteria. | 5-year recurrence rate 12.5% |
| South Korea; Choi et al., 2021 (Clin Mol Hepatol)  PMID: 33525077 | 2014–2017 | 2,563 | Multiple criteria. | HCC recurrence rate 6.7% (LDLT) and 6.7% (DDLT) |

**Supp. Tab. 1. Contemporary multicenter cohorts reporting HCC recurrence after liver transplantation.** Abbreviations: AC-DS, all-comers downstaging; AFP, alpha-fetoprotein; CI, confidence interval; DDLT, deceased-donor liver transplantation; DS, downstaging; HCC, hepatocellular carcinoma; LDLT, living-donor liver transplantation; LRT, locoregional therapy; LT, liver transplantation; PMID, PubMed identifier; UNOS, United Network for Organ Sharing; UNOS-DS, UNOS downstaging criteria.
