## Supplementary Materials for "Engineering CAR T Cells for Hepatocellular Carcinoma Recurrence after Liver Transplantation"

Animal studies

NOD-*scid* IL-2Rg^null^ (NSG) mice (NCI Frederick) were housed and maintained under a protocol approved by the NIH Institutional Animal Care and Use Committee. All mice were used at 6-12 weeks old unless otherwise noted. For the peritoneal Hep G2 model, 2 x 10^6^ luciferase-expressing Hep G2 (Hep G2-Luc) cells were injected intraperitoneally (i.p.) as previously described (11). Two weeks after tumor inoculation, mice were randomized to achieve comparable baseline tumor burden. Mice received i.p. injections of 2 x 10^7^ CAR T cells or untransduced T cells. Prior to cell administration, mice were pretreated with 1 mg/kg tacrolimus i.p. or an equivalent volume of DMSO vehicle control. Tacrolimus and control treatment was continued at 0.5 mg/kg twice daily for four additional days. Tumor burden was assessed by total bioluminescent flux using an IVIS Lumina imaging system (PerkinElmer) 7 min after i.p. injection of 3 mg D-Luciferin (Syd Lab, #MB000102-R70170) by a blinded investigator. Photon flux of the entire tumor tissue was calculated using the Living Image software 4.8.4 (IVIS™ imaging systems). All experiments were conducted according to local institution guidelines and approved by the Animal Care and Use Committee of the National Institutes of Health (NIH), Bethesda, USA.

To assess the alloreactive potential of *FKBP1A*/*TRAC* double-knockout CAR T cells, NSG mice were used in a xenogeneic graft-versus-host disease model. Mice were sublethally irradiated with 1.3 Gy and subsequently injected intravenously with 1 × 10⁷ wild-type CAR T cells or CD3-depleted *FKBP1A*/*TRAC* double-knockout CAR T cells. Animals were monitored longitudinally for clinical signs of GvHD and euthanized according to predefined humane endpoint criteria. Peripheral blood was collected at serial time points to quantify circulating human T cells by flow cytometry, defined as human CD45-positive cells among total human and murine CD45-positive leukocytes, and to determine the frequency of residual TCR/CD3-positive cells within the human T cell compartment. At euthanasia or at the experimental endpoint, spleens and livers were harvested for flow cytometric and histopathological analysis. Splenocytes were analyzed for human T cell engraftment and residual endogenous TCR/CD3 expression, while liver tissues were fixed, sectioned, and stained with hematoxylin and eosin to assess immune-mediated tissue injury.

Cell lines

HepG2 (hepatoblastoma) and Hep3B (hepatoma) cell lines expressing GFP and firefly luciferase were kindly provided by Mitchell Ho. The cell lines were routinely tested for mycoplasma contamination. Patient PBMCs were collected from patients 30 days after allogeneic liver transplantation. Healthy control PBMCs were provided by the NIH Blood Bank.

CAR T cell production

Peripheral blood mononuclear cells (PBMCs) from healthy volunteers were isolated using Lympholyte cell separation medium (Thomas Scientific, #CHM03X128), followed by untouched Pan T cell isolation (Miltenyi, #130-096-535). T cells were rested overnight in T cell medium (RPMI 1640 supplemented with 10% FCS, 1% penicillin/streptomycin, 1% Na-pyruvate, 0.1% HEPES, 1% non-essential amino acids, and 50 µM β-mercaptoethanol) supplemented with human IL-7 (5 ng/mL). The following day, cells were washed in PBS prior to electroporation. sgRNAs containing 2′-O-methyl modifications at the first and last three bases and 3′ phosphorothioate linkages between the first three and last two bases were purchased from Synthego. A total of 15 µg *TRAC* locus targeting sgRNA and 15 µg of *FKBP1A* locus targeting sgRNA were mixed at a 1:1 ratio and incubated with 30 µg recombinant S. pyogenes high fidelity Cas9 nuclease (Integrated DNA Technologies, #1081061) for 20 minutes to form ribonucleoprotein complexes. Between 1 x 10^6^ and 1 x 10^7^ cells were electroporated using the EO-115 program on the Lonza 4D-Nucleofector X Unit with the P3 Primary Cell 4D-Nucleofector X Kit L, according to the manufacturer’s instructions. Electroporated T cells were transferred into RPMI prewarmed to 37 °C and incubated for 5 minutes, followed by the addition of 2x T cell medium supplemented with recombinant human IL-2 (200 IU/mL, PeproTech), IL-7 (10 ng/mL, PeproTech), and IL-15 (10 ng/mL, PeproTech). T cells were stimulated at a 1:2 bead-to-cell ratio using CD3/CD28 (Thermofisher, #11452D) beads or with 50 µl TransAct (Miltenyi, #130-128-758) per 5 x 10^6^ cells. After 48 hours, T cells were transduced with a second-generation lentiviral vector encoding an anti-GPC3 CAR, as previously described (11). Cells were then expanded for additional 5 to 8 days before use in functional *in vitro* and *in vivo* assays. Transduction efficiency was assessed by flow cytometry based on staining for a truncated EGFR (EGFRt) tag expressed downstream of a 2A sequence in the CAR construct. To ensure comparability, transduction efficiencies were normalized between electroporated and non-electroporated cells by adding untransduced T cells.

Killing assay

Tumor cell cytotoxicity was evaluated using luciferase-expressing HepG2 and Hep3B target cells. Tumor cells were seeded in 96-well flat-bottom plates and allowed to adhere overnight. CAR T cells or untransduced T cells were added at the indicated effector-to-target (E:T) ratios in the presence or absence of tacrolimus (15 ng/mL) or cyclosporine A (300 ng/mL). After 24 to 48 hours of co-culture, residual viable tumor cells were quantified by bioluminescence measurement following addition of D-luciferin substrate according to the manufacturer's instructions (Promega). Luminescence was measured using a plate reader, and specific lysis was calculated relative to tumor-only control wells. For cytokine analyses, supernatants were collected before luciferase measurement and stored at -20°C until analysis.

Antigen-specific assays

Ninety-six-well plates were coated overnight at 4°C with recombinant human GPC3-IgG1 Fc fusion protein (5 µg/mL in PBS; AcroBiosystems, #GP3-H5258). The following day, the coating solution was discarded, and 1 × 10^5^ untransduced T cells or CAR T cells were plated in RPMI medium and cultured for 48 hours. IFN-γ concentrations in culture supernatants were quantified using a human IFN-γ ELISA kit according to the manufacturer's instructions (BD Biosciences, #555142). Absorbance was measured at 450 nm using a microplate reader, and cytokine concentrations were calculated from a standard curve generated using recombinant IFN-γ.

In vitro stimulation assay

To assess resistance to immunosuppressive agents, T cells were stimulated with phorbol 12-myristate 13-acetate and ionomycin in the presence of brefeldin A for 4 hours at 37°C. Cells were cultured with vehicle control or tacrolimus at the indicated concentrations. Following stimulation, cells were stained with viability dye and antibodies against surface markers, fixed and permeabilized using the Cytofix/Cytoperm kit (BD Biosciences, #554714), and stained intracellularly for IFN-γ. Samples were analyzed by flow cytometry and data were processed using FlowJo software (BD Biosciences).

Knockout selection

To enrich gene-edited T cells, pharmacologic selection was initiated 48 hours after electroporation and T cell activation. Everolimus, sirolimus, or tacrolimus was added to cultures at 100 ng/mL and maintained throughout the expansion period. Culture medium and cytokines were replenished every two days. At the end of the culture period, cells were harvested and analyzed for TCR expression by flow cytometry and for editing efficiency by genomic sequencing. Cell expansion, viability, and phenotype were assessed to determine the impact of each selection strategy on product composition. For combined enrichment approaches, cultures were first subjected to mTOR inhibitor-mediated selection using everolimus or sirolimus, followed by depletion of residual CD3-positive cells using magnetic-activated cell sorting (MACS) prior to downstream functional analyses.

Genomic DNA isolation and editing analysis

Genomic DNA was isolated from edited T cells using the QIAamp DNA Blood Mini Kit (Qiagen, #51106) according to the manufacturer's instructions.

Sanger sequencing. Genomic regions flanking the FKBP1A and TRAC sgRNA target sites were amplified by PCR using locus-specific primers and Q5 High-Fidelity Polymerase (NEB, #M0491L) under the following cycling conditions: 50 ng DNA, 98°C for 10 s, 68°C for 20 s, and 72°C for 20 s for 30 cycles. PCR products were purified using the DNA Clean & Concentrator kit (Zymo Research, #D4030) and subjected to Sanger sequencing. Editing efficiencies were quantified from sequencing chromatograms using the Tracking of Indels by DEcomposition (TIDE) algorithm.

Next-generation sequencing (NGS). For each sample, 10 ng of genomic DNA was used as template to amplify regions surrounding the TRAC and FKBP1A sgRNA target sites using the primers listed below. PCR reactions were performed in 10 μL volumes using Q5 Ultra II 2× Master Mix (NEB, #M0544X) under the following cycling conditions: 98°C for 15 s, 68°C for 15 s, and 72°C for 15 s for 30 cycles. A second PCR was then performed to add the remaining Illumina adapter sequences and unique sample barcodes. Libraries were sequenced on an Illumina MiSeq platform using 2 × 150 bp paired-end sequencing. Raw FASTQ files were processed using a previously described amplicon sequencing analysis pipeline (<https://github.com/rajchari2/ngs_amplicon_analysis>) to quantify editing frequencies at each target site (PMID: 33051688).
